# Unraveling the Genetic Complexity of Invasive *Lonicera* spp.: Evidence of Hybridization from Nuclear and Chloroplast Genome Analysis

**DOI:** 10.1101/2024.12.25.630324

**Authors:** Michael V. Osier, Eli J. Borrego, Samantha V. Tran, André O. Hudson

## Abstract

Members of the *Lonicera* genus, commonly known as honeysuckles, encompass both native and invasive species in North America, the latter posing significant ecological challenges. Invasive honeysuckles can disrupt native ecosystems by outcompeting local flora, altering habitats, and affecting wildlife populations. Effective management and control of these species are critical but hindered by the difficulty of accurate identification due to overlapping morphological traits. Traditional control measures recommended by government agencies, such as cutting, burning, and chemical treatments, have proven only partially effective, sometimes complicated by misidentification of species.

The morphological identification of invasive honeysuckles is notoriously challenging. Key distinguishing features are often subtle, variable, and sometimes only discernible during specific developmental stages, complicating field identification efforts. This taxonomic ambiguity underscores the urgent need for more precise identification methods that transcend conventional morphological assessments.

In this study, we explore the molecular identity of a Lonicera isolate collected from the Finger Lakes region of Western New York. Through detailed genetic analysis, we demonstrate that the specimen likely possesses maternal inheritance from *Lonicera tatarica* while the majority of its nuclear genome is associated with *Lonicera insulara*. Both species are prevalent in the region and known for their invasive potential.

Our findings highlight the utility of molecular techniques as a complementary tool for species identification within the Lonicera genus. By providing a clearer genetic framework for distinguishing between morphologically similar species, this approach can enhance conservation strategies, inform management decisions, and improve ecological restoration efforts. As invasive species continue to threaten biodiversity, integrating molecular diagnostics with traditional methods offers a promising path toward more effective environmental stewardship.

## Introduction

In the eastern United States, several *Lonicera* species originating from Asia and Europe have become invasive shrubs, raising concerns for state environmental conservation departments. These species were initially introduced to the United States as ornamental shrubs for landscaping due to their attractive appearance and hardiness. However, their ecological impact has become problematic over time. Members of the Tataricae clade of invasive *Lonicera* are particularly concerning because of their ability to grow as upright, multi-stemmed shrubs with expansive canopies that can reach several meters in both height and width. Their dense foliage creates heavy shading that suppresses the growth of hardwood seedlings, ultimately hindering forest regeneration and altering natural successional processes [1]. Furthermore, these invasive shrubs have been linked to reduced species richness in the areas they colonize [1–4].

The morphological identification of the Tataricae clade within the Coeloxylosteum botanical section, and *Lonicera* species more broadly, is notoriously challenging due to significant overlap in leaf and stem characteristics among species. Leaf shapes are notably similar across species and even change throughout the life cycle of individual leaves. For example, in *Lonicera insularis*, which has been synonymized with *Lonicera morrowii* by Srivastav et al. [5], leaves exhibit dynamic morphological changes that complicate identification. Even more distantly related species, such as *Lonicera maackii*, have leaves that are oblong but pointed at the tips, a trait also observed in certain stages of *L. insularis*. All these species share common features such as hollow, woody stems and pairs of red berries arranged oppositely along the stem. Flower color offers some distinction, with *L. insularis* typically producing white flowers, while *Lonicera tatarica* flowers range from pink to purple.

However, hybridization further complicates morphological identification. As noted by Barnes and Cottam [6], hybrids *of L. insularis* and *L. tatarica*, collectively referred to as *Lonicera × bella*, are widespread throughout the northeastern and midwestern United States. These hybrids exhibit continuous phenotypic variation between the parental species [6]. In some regions, hybrid populations may show traits skewed toward one parent, while others display a full spectrum of intermediate characteristics. This frequent hybridization, combined with limited distinguishing morphological features, underscores the difficulty of relying solely on physical traits for species identification. Consequently, alternative methods such as genetic analysis could significantly enhance conservation efforts.

Previously, we published the nuclear genome sequence of what was initially identified morphologically as *L. maackii* (NCBI PRJNA521295 [7]). At the time, limited genetic reference data were available, constraining our ability to confirm species identity through molecular methods. More recently, Srivastav et al. (2023) released RADSeq data for 72 *Lonicera* species (PRJNA925902 [5]), providing an essential comparative resource.

In this study, we leveraged the RADSeq dataset from Srivastav et al. (2023) to perform molecular species identification of our previously sequenced *Lonicera* specimen. Additionally, we isolated chloroplast genome (cpDNA) sequences from our 2022 *Lonicera* sample and compared them to cpDNA sequences from members of the Coeloxylosteum clade. By integrating nuclear and chloroplast genetic data, we determined that our specimen most likely inherited its maternal lineage from *L. tatarica*, while its nuclear genome shows primary inheritance from *L. insularis*. We also acknowledge the possibility of additional admixture with other *Lonicera* species. This study represents the first molecular evidence confirming hybridization within *Lonicera* populations in natural habitats and highlights the utility of combining nuclear RADSeq and cpDNA sequences for accurate species identification in hybrid complexes across the genus.

## Materials and Methods

### Nuclear genome analysis

The specimen identification, DNA extraction, sequencing methodology, and assembly are detailed in our previous study [7]. Said nuclear genome sequence was used to create a BLAST nucleotide database using BLAST+ ver. 2.15 [8]. The first 100,000 sequence fragments for each available *Lonicera* spp. were selected from the raw RADSeq data [5] and extracted as FASTA files. These sequences were then further divided into samples containing one of: the full-length sequence, the first 40 nucleotides, or the first 30 nucleotides. These sample files were then used to query the BLAST database of our specimen searching for an ungapped match with no mismatched nucleotides. The percentage of RADSeq fragments from each species and fragment length that had a full-length match to our specimen database was determined. Estimated heterozygosity was calculated by back-mapping reads to the genome assembly using the STAR aligner version 2.5.4b [9], and calling variation with samtools ver. 1.6 [10] and bcftools ver. 1.15.1-14 [10] to estimate observed variation in the specimen, then dividing by the assembly length.

### cpDNA identification

Raw sequence data from our specimen was cleaned using FastQC ver. 0.12.1 (https://www.bioinformatics.babraham.ac.uk/projects/fastqc/) and FastX ver. 0.0.13 (https://github.com/agordon/fastx_toolkit) to remove low quality reads, then aligned to the *L. maackii* cpDNA sequence (MN256451.1 [11]) and *L. tatarica var morrowii* (OQ784205.1) using the STAR aligner version 2.5.4b [9]. The consensus sequence from each alignment was extracted to a FASTA file with samtools [10] and again used to create a BLAST nucleotide database using BLAST+ ver 2.15 [8]. Published cpDNA sequence of the coding regions for *L. insularis* (NC_039634.1), *L. tatarica* (OQ784187.1), *L. tatarica var morrowii* (OQ784205.1, presumably the same as *Lonicera x bella*), and *L. maackii* (NC_039636.1) were then used to query the database of our specimen with BLAST+ ver. 2.15 [8] with a minimum of 80% query coverage and only saving the best hit. cpDNA sequences above and from additional species in the Isika clade (*L. discolor*, OQ784177.1; *L. maximowiczii*, NC_050941.1; *L modesta var. lushanensis*, OM141002.1; *L nervosa*, NC_040961.1; *L ruprechtiana*, OQ784189.1 and NC_056986.1; *L. xylosteum*, OQ784191.1), two additional *L. maackii* cpDNA sequences (MW784237.1 and MN256451.1), and one additional *L. tatarica* (OQ784187.1) were aligned with MAFFT version 7.505 [12]. A best-fit model consensus tree was generated using IQ-TREE [13,14].

## Results

### Nuclear genome identification

The counts of alignment matches between random RADSeq fragments displayed expectations of a normal distribution across species, with a skewed tail toward species with higher match rates (Fig 1). Among the species with the most matches, seven were at an inflection point above the distribution (Table 1). All but two of these best matching species, belong to Tataricae [5]. Note that *L. tatarica var morrowii* is the hybrid of *L. insularis* (also known as *L. morrowii*) and *L. tatarica*, and had a number of matches intermediate between its two parental species (Table 1). *L. sovetkinae* and *L. korolkowii* are both more distantly related, potentially reflecting a lack of direct ancestry with the founding invasive plants.

**Table 1.** Number of k-mer alignment matches for the top *Lonicera* spp. of Figure 1. Species marked in bold belong to the Tataricae clade of *Lonicera*.

|  | Matches by k-mer length |  |  |
| --- | --- | --- | --- |
| <u>Species</u> | <u>30-mer</u> | <u>40-mer</u> | <u>Full-length</u> |
| ***L. insularis*** | 72943 | 39465 | 33839 |
| ***L. tatarica var morrowii*** | 60197 | 34783 | 29354 |
| ***L. tatarica*** | 51420 | 24586 | 17562 |
| *L. quinquelocularis* | 43658 | 12050 | 10267 |
| ***L. sovetkinae*** | 38606 | 25758 | 19878 |
| *L. obovata* | 38382 | 6028 | 5133 |
| ***L. korolkowii*** | 37750 | 24028 | 20517 |
| *L. maackii* | 14442 | 5190 | 4198 |

**Fig 1.**
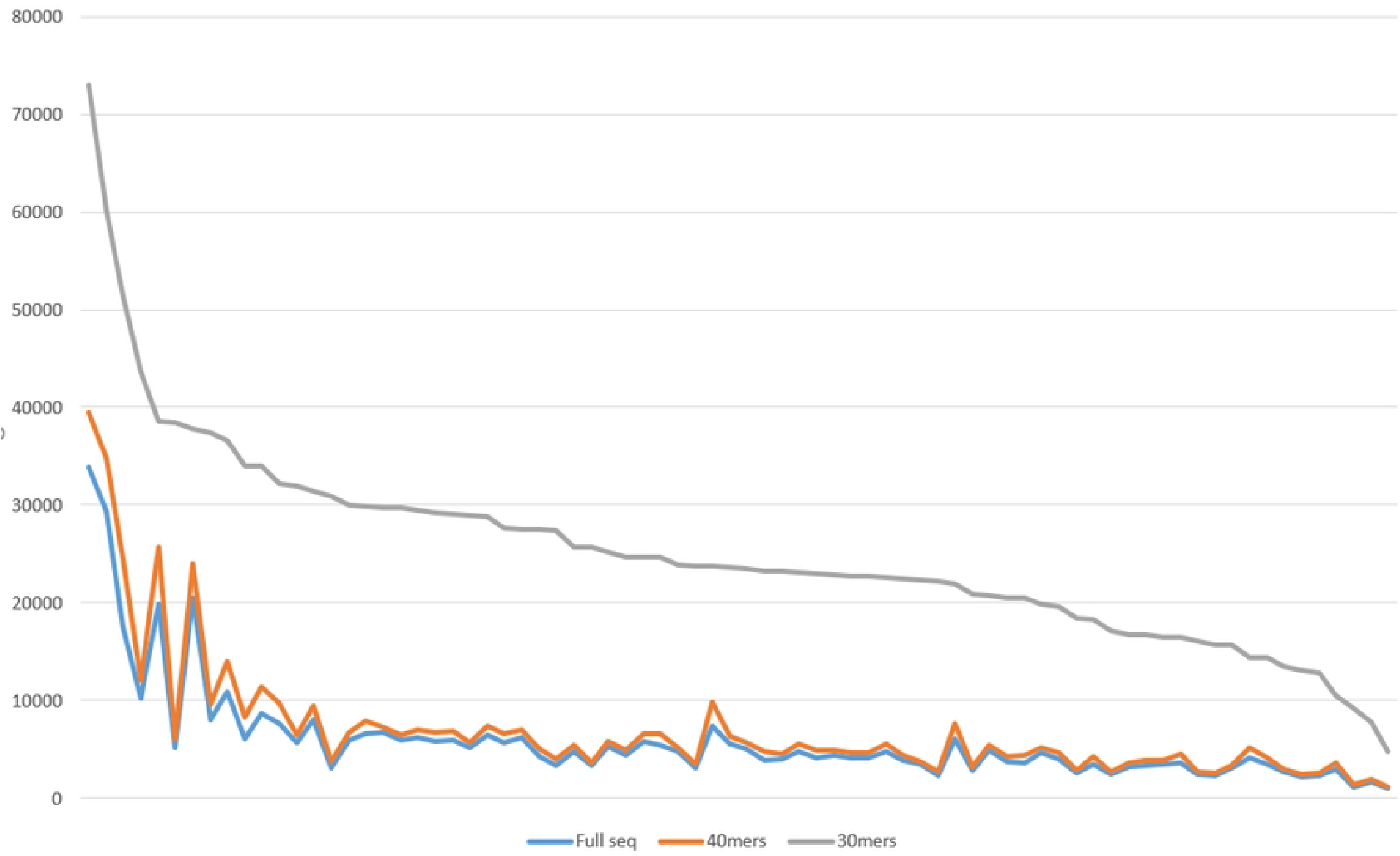
Distribution of the number of sequence alignment matches. The x-axis represents species, and the y-axis represents counts of RADSeq fragments with exact matches to the Kesel genome. Line colors indicate fragment size used for comparison.

The observed heterozygosity of the sequenced specimen was 0.4%. At this heterozygosity, one would expect 88.7% of 30-mers to match exactly, or 88,670 BLAST hits. We observed 72,943 BLAST hits for the best matching species. Although this value approaches the expectation, the observed matches are slightly lower. This leaves open the possibility that there is additional species admixture, possibly from one of the other closely matching species.

### cpDNA genome identification

Hybrid *Lonicera* are reported across the United States and anticipated in the Finger Lakes region based on mixed or intermediate morphology in local populations. Since our specimen repeatedly mapped to members of the Tataricae clade, we compared the coding regions of its cpDNA against that of *L. insularis* and *L. tatarica* for confirmation and to investigate the possibility of hybridization. We also compared it against *L. maackii* to test that the specimen does not belong to that species. 85 genes/coding regions were tested. With the exception of 15 genes (psbA, psbI, petN, psbM, psbZ, rps4, atpE, petA, psbL, petL, petG, rpl33, rpoA, ycf15, ndhE), which showed perfect identity across all three genomes at the nucleotide level, all but two *L. maackii* matches had a lower percent identity than either *L. insularis* and *L. tatarica*, providing strong molecular evidence that the specimen does not belong to *L. maackii* (Table 2).

**Table 2.**
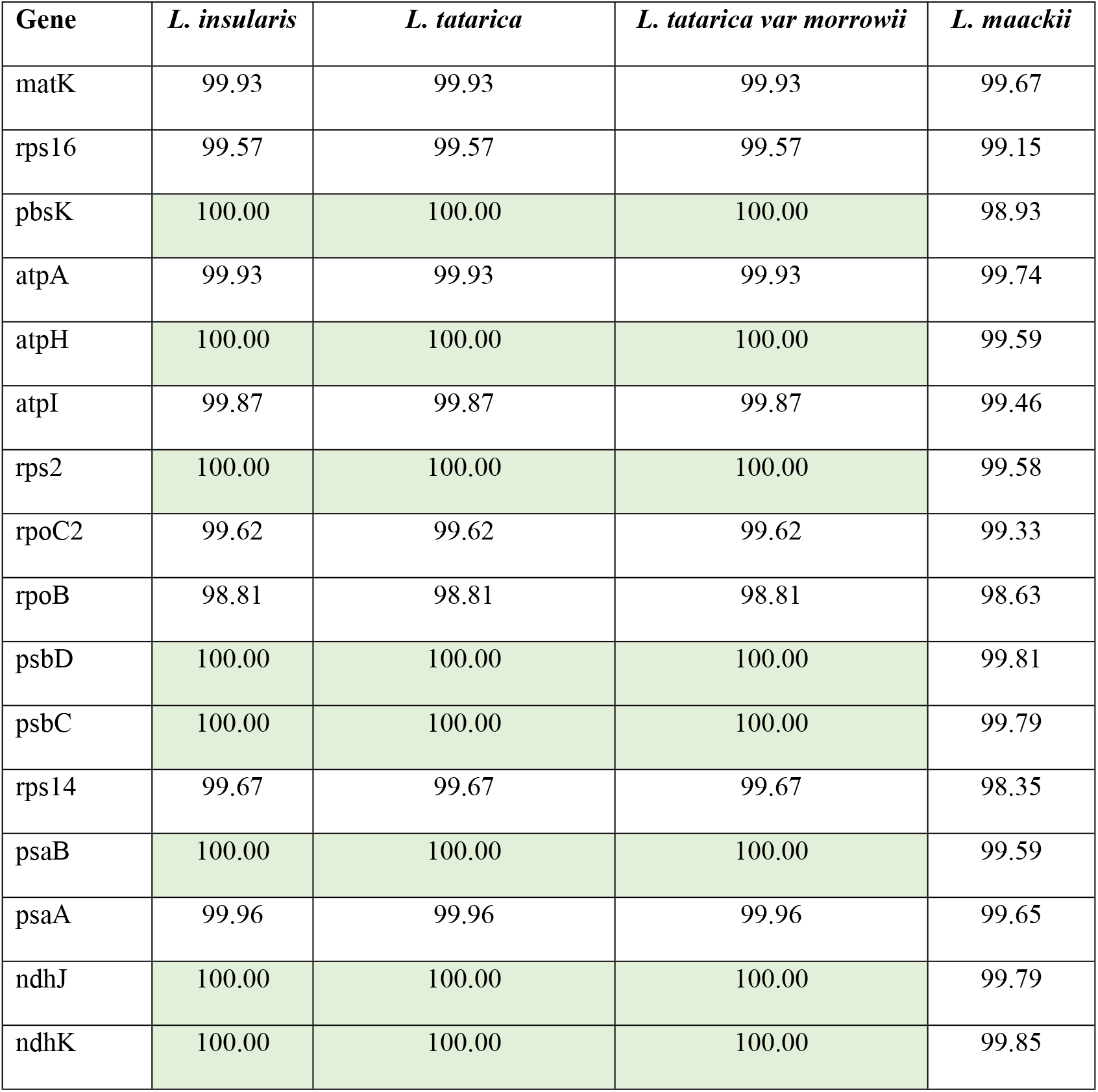
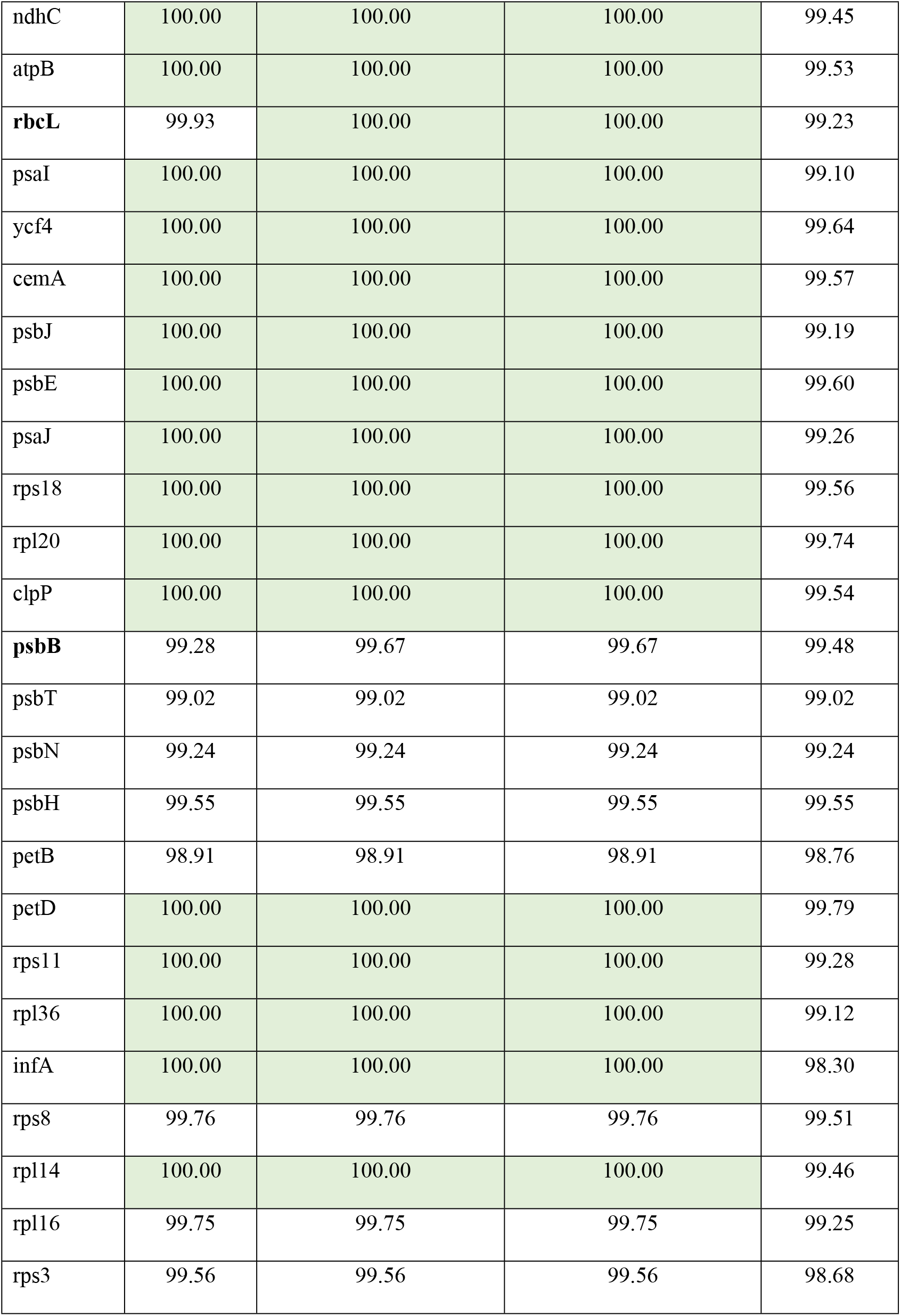
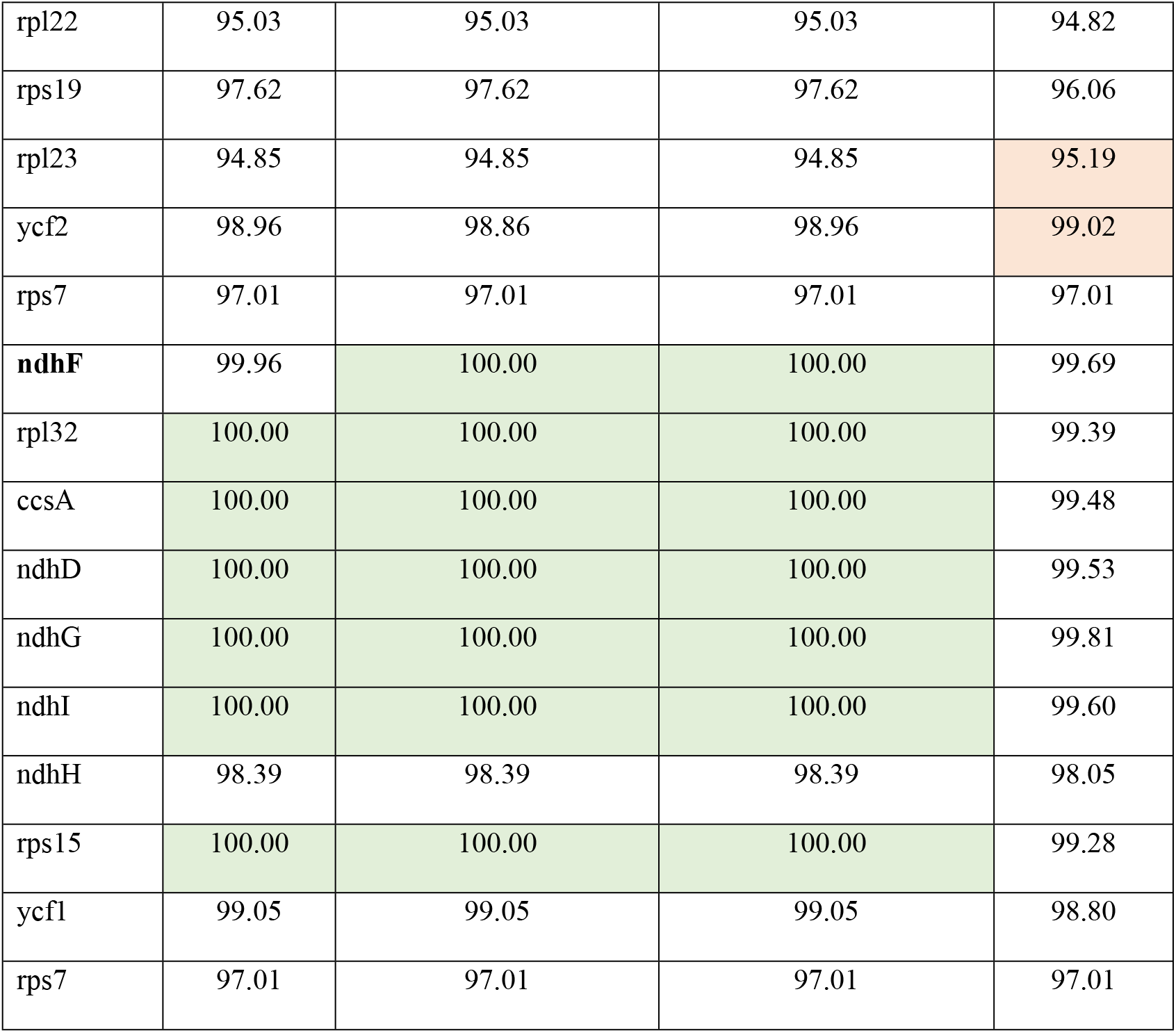
Comparison of selected chloroplast genes between our specimen and *L. insularis, L. tatarica*, and *L. maackii*. Percent identities of alignments between our specimen and *Lonicera spp*. for selected chloroplast coding regions of *Lonicera spp*. Identities of 100% are highlighted in green and genes with perfect identity across all three species were removed. Genes where *L. tatarica* is closer to the specimen are in bold. Genes were *L. maackii* displayed the highest identity in comparisons to our specimen are highlighted in red.

The percent identity of the nucleotide sequence of genes were nearly all the same between our specimen and either *L. insularis* or *L. tatarica var morrowii*, with four exceptions. For rbcL, there was a single synonymous G→A nucleotide difference where our sample matched *L. tatarica var morrowii* and *L. tatarica*. Other differences between *L. insularis* and both our sample and *L. tatarica var morrowii* were six nucleotides of psbB and one nucleotide of ndhF. The last gene that showed difference contained a single C in *L. insularis* and a T in *L. tatarica* and *L. tatarica var morrowii*.

### cpDNA phylogeny in Coeloxylosteum

To better understand the chloroplast genome ancestry of the specimen, entire cpDNA genome sequences were phylogenetically compared among members of Coeloxylosteum, as well as members of the botanical section Isika, a closely related outgroup for Tataricae. Complete cpDNA genome sequences were not available for two out the four members of Tataricae: *L. sovetkinae* and *L. korolkowii*. To ensure that the method of aligning to a reference did not introduce bias in favor of the reference species, specimen data was aligned to both *L. maackii* and *L. tatarica var morrowii* reference cpDNA. Both consensus alignments of the Kesel data were included in the phylogenetic analysis, and clustered together (Fig 2) and close to, but slightly outside, the clade of Tataricae species. However, the specimen was distinct from *L. maackii*, as per the nuclear genome observations.

**Fig 2.**
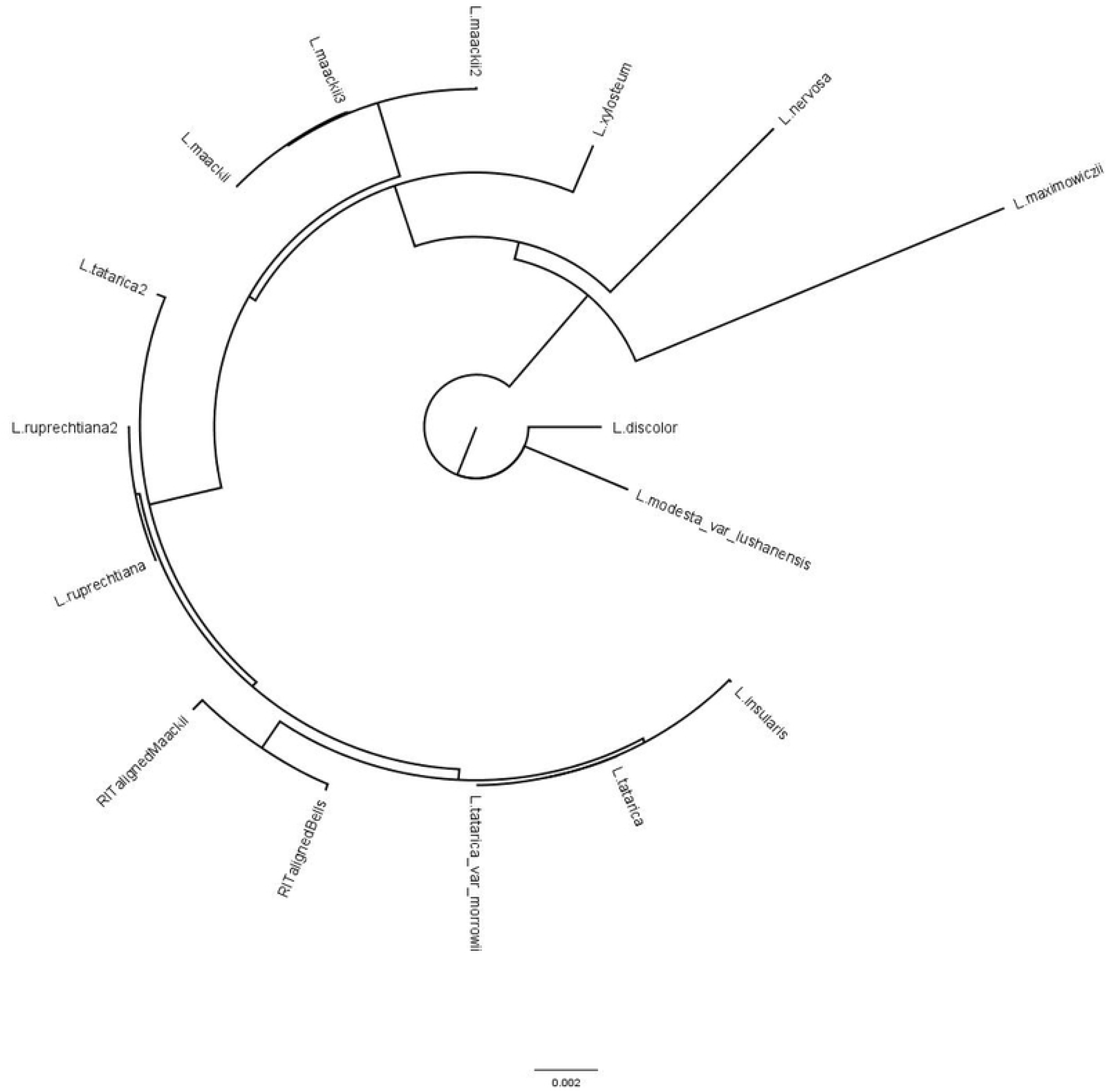
Phylogenetic analysis of the specimen in reference to the Tataricae and Isika members, as well as the *L. maackii* outgroup.

## Discussion

Although initially supported by morphological characteristics, the differences observed in the nuclear genome sequences strongly suggest that our previously published *Lonicera* sp. [7] is almost certainly not *L. maackii*. Despite this, no species among the 72 *Lonicera spp*. tested showed sufficiently high percentage of mapped reads to our published genome, making the definitive identification of our specimen as a single species unlikely.

The k-mer distribution from our prior study (Supplementary Figure S1 [7]), interpreted using Merqury [15], revealed a pronounced second peak, indicating a low level of heterozygosity in the specimen. This finding was supported by the low genetic variation observed. It is possible that our specimen represents a highly inbred lineage, which would artificially reduce heterozygosity even if the regional population were genetically diverse. In such a scenario, the observed lower match rate between *L. insularis* and our specimen would be expected. Resolving this issue would require a population-level study to assess the heterozygosity of local *Lonicera* populations.

In our cpDNA analysis, we considered the potential for sequence bias due to aligning reads to *L. maackii* and *L. tatarica var. morrowii*. While the STAR aligner allows for a degree of sequence divergence reducing alignment bias, the selection process could nonetheless preferentially favored sequences similar to the previously published *L. maackii* cpDNA. However, nearly all coding sequences from other *Lonicera spp*. were identified, often with complete coding regions, suggesting minimal alignment bias. Notably, all cpDNA gene sequence differences identified matched *L. tatarica*, and no nucleotides specific to *L. insularis* were detected. If our specimen were indeed *L*. insularis, as indicated by the nuclear genome, some *L. insularis*-specific cpDNA sequences should have been present.

Interestingly, a small number of bases in the cpDNA coding regions showed greater similarity to *L. maackii* than to the Tataricae clade species, and additional intergenic differences were observed that did not match any known species. These findings raise the possibility that our specimen has accumulated unique cpDNA mutations since its introduction to the U.S. over 150 years ago, or that its cpDNA originates from an unidentified species. Sequences for *L. sovetkinae* and *L. korolkowii* were unavailable, leaving open the possibility of their maternal contribution, although no records have yet document their presence as invasive species in the Finger Lakes region of New York.

The discrepancy between nuclear and plastid genome species identifications supports the hypothesis of hybridization of *Lonicera* spp. in this region, consistent with previous findings in the Midwestern U.S. [6]. Given the limited sequence divergence in the cpDNA of the Tataricae clade, and the lack of perfect identity to our specimen, additional possibilities for maternal inheritance remain. While it is unlikely that our specimen represents a new species due to the extensive literature on *Lonicera* taxonomy, regional genetic differentiation within founder populations introduced during the 19th century could explain the observed cpDNA variation. Founder-related bottlenecks, early mutations after introduction, or regionally distinct variants could all contribute to these genetic differences.

Alternatively, our specimen could be a recent transplant from a genetically distinct population in Asia or Europe not represented in current reference sequences. A comprehensive population genetics study involving both nuclear and plastid genomes would be essential to disentangle these possibilities and clarify the evolutionary history of this complex species group.

## Conclusion

This study provides molecular evidence supporting the hybrid nature of our *Lonicera* specimen from the Finger Lakes region of New York. Through a comprehensive analysis of nuclear and chloroplast genome sequences, we identified a likely maternal contribution from *L. tatarica* and substantial nuclear genome inheritance from *L. insularis*. The observed discrepancies between nuclear and plastid genome identifications reinforce the hypothesis of interspecific hybridization, a phenomenon previously documented in the genus but as of yet not characterized at the molecular level.

The results also underscore the challenges associated with species identification in the *Lonicera* genus, where overlapping morphological traits and frequent hybridization complicate traditional taxonomic approaches. Our findings highlight the necessity of integrating molecular techniques such as RADSeq and cpDNA analysis into conservation and management strategies for invasive *Lonicera* populations. These approaches provide more precise species identification, guiding better-informed ecological interventions and restoration efforts.

Furthermore, the possibility of region-specific genetic variation due to historical introduction, founder effects, or undocumented transplants points to the need for broader population-level studies. A comprehensive population genetics approach could reveal the full extent of genetic diversity within invasive *Lonicera* populations and inform efforts to mitigate their ecological impact.

In conclusion, our molecular analysis advances the understanding of *Lonicera* hybridization dynamics and demonstrates the utility of genomic tools in addressing taxonomic uncertainties. By applying these methods, future studies can enhance conservation strategies for managing invasive honeysuckle species and protecting native ecosystems from their detrimental effects.

## Acknowledgements

The authors thank the reviewers for their time and providing cogent feedback. The authors thank the Thomas H. Gosnell School of Life Sciences and the College of Sciences at the Rochester Institute of Technology for ongoing support.

